# Fluence-dependent degradation of fibrillar type I collagen by 222 nm far-UVC radiation

**DOI:** 10.1101/2023.09.19.558392

**Authors:** Antonia Kowalewski, Nancy R Forde

**Affiliations:** Department of Physics, Simon Fraser University, Burnaby, BC, V5A 1S6, Canada; Department of Molecular Biology and Biochemistry, Simon Fraser University, Burnaby, BC, V5A 1S6, Canada

## Abstract

For more than 100 years, germicidal lamps emitting 254 nm ultraviolet (UV) radiation have been used for drinking-water disinfection and surface sterilization. However, due to the carcinogenic nature of 254 nm UV, these lamps have been unable to be used for clinical procedures such as wound or surgical site sterilization. Recently, technical advances have facilitated a new generation of germicidal lamp whose emissions centre at 222 nm. These novel 222 nm lamps have commensurate antimicrobial properties to 254 nm lamps while producing few short- or long-term health effects in humans upon external skin exposure. However, to realize the full clinical potential of 222 nm UV, its safety upon internal tissue exposure must also be considered. Type I collagen is the most abundant structural protein in the body, where it self-assembles into fibrils which play a crucial role in connective tissue structure and function. In this work, we investigate the effect of 222 nm UV radiation on type I collagen fibrils *in vitro*. We show that collagen’s response to irradiation with 222 nm UV is fluence-dependent, ranging from no detectable fibril damage at low fluences to complete fibril degradation and polypeptide chain scission at high fluences. However, we also show that fibril degradation is significantly attenuated by increasing collagen sample thickness. Given the low fluence threshold for bacterial inactivation and the macroscopic thickness of collagenous tissues *in vivo*, our results suggest a range of 222 nm UV fluences which may inactivate pathogenic bacteria without causing significant damage to fibrillar collagen. This presents an initial step toward the validation of 222 nm UV radiation for internal tissue disinfection.

## Introduction

Surgical-site infections are a leading cause of health care-associated infections worldwide, affecting up to one-third of surgical patients in low- and middle-income countries and up to 10% of surgical patients in Europe [1]. The majority of these infections are caused by antimicrobial-resistant pathogens [1]. Ultraviolet C (UVC) radiation, defined as electromagnetic radiation of wavelengths 200 to 280 nm, effectively inactivates both drug-sensitive and drug-resistant bacteria [2–11]. While this property has been exploited for water and surface disinfection for over 100 years, the use of UVC radiation for medical purposes has remained limited due to the mutagenic effects of 254 nm UV, the most readily available UVC wavelength [12, 13]. More recently, however, technical advances have facilitated the production of shorter-wavelength 222 nm “far-UVC” lamps which have commensurate anti-bacterial properties to 254 nm UV lamps but do not cause mutagenic DNA damage upon external human skin exposure [2, 14, 15]. Given its high level of absorbance by proteins, it is proposed that far-UVC radiation is absorbed by the protein-rich anucleate cells of the stratum corneum before it can reach the nucleated cells of the lower epidermis and induce DNA damage [2]. Some studies suggest that far-UVC radiation is also unable to penetrate the cytoplasm of individual unprotected eukaryotic cells [16, 17]. In bacterial cells, however, due to a small cell volume and lack of a protective stratum corneum, far-UVC radiation remains cytotoxic [2].

So far, investigations into the human safety of far-UVC radiation have focused on external skin exposure [2, 15, 18–22]. Despite this, far-UVC radiation is actively being proposed for open-wound sterilization and surgical-site disinfection [6, 14, 19, 23, 24]. To determine the safety of far-UVC radiation for such exposures, it is necessary to investigate its effect on the body’s internal tissues. In this study, we investigate the effect of 222 nm far-UVC radiation on type I collagen fibrils, the main protein component of the extracellular matrix (ECM) and connective tissues including tendons, ligaments, skin, bone, and cartilage [25, 26].

Type I collagen fibrils are composed of laterally associated type I collagen molecules, each of which is a heterotrimer of two α_1_ and one α_2_ polypeptide chains arranged in a 300 nm-long right-handed triple helix [27, 28] (Fig. 1). Fully formed fibrils range from 30 to 300 nm in diameter and are stabilized by non-covalent interactions between collagen molecules [29–31]. In the ECM, collagen fibrils form a network which can be modelled experimentally by a fibrillar gel [32] (Fig. 1).

**Fig 1.**
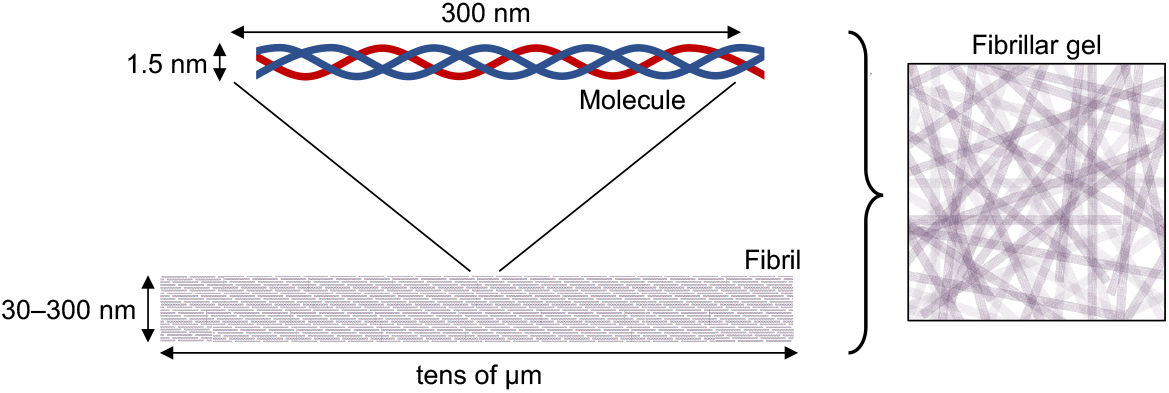
Structure of a type I collagen fibrillar gel. Blue lines represent α_1_ chains and red lines represent α_2_ chains.

To the authors’ knowledge, this study presents the first investigation into the effect of 222 nm far-UVC radiation on type I collagen.

## Materials and methods

### Sample preparation

Collagen fibrils were formed by combining rat tail tendon-derived acid-soluble type I collagen (5 mg/mL in 20 mM acetic acid) (CultrexRat Collagen I, R&D Systems) with 400 mM HEPES + 0.02% sodium azide and 1 M NaCl at room temperature to reach final concentrations of 1 mg/mL collagen, 100 mM HEPES + 0.005% sodium azide, and 270 mM NaCl. Samples were added to a clear-bottom 96-well plate (Nunc MicroWell 96-Well Optical-Bottom Plate with Polymer Base, Thermo Fisher Scientific) (60 µL per well) or flowed into a 0.175 or 1.2 mm-thick microscope sample chamber and sealed with nail polish. Sample chambers consist of a 1.0 mm–thick ge124-grade quartz microscope slide (75875-470, VWR) and a 0.13 to 0.16 mm–thick borosilicate glass coverslip (12-542B, Fisherbrand) separated by two parallel stacks of double-sided tape (Tesa) spaced approximately 5 mm apart. All fibrillar collagen samples were left to polymerize at room temperature for at least four hours prior to irradiation and analysis. Fibril formation under these conditions was confirmed by monitoring the optical density of the solution at 347 nm with a BioTek Synergy HTX microplate reader (Agilent) with 100 µL per well (Fig. Figure S1). Molecular collagen samples were prepared in the same way as fibrillar collagen samples (1 mg/mL in 100 mM HEPES + 0.005% sodium azide and 270 mM NaCl) but were irradiated immediately following neutralization, before fibril formation could occur.

Bovine serum albumin (BSA) samples were prepared by diluting lyophilized BSA powder (Sigma Aldrich) to 1 mg/mL in ddH_2_O and adding to a clear-bottom 96-well plate (60 µL per well).

### UV irradiation

Sample irradiation was performed using a LILY Handheld Personal Far UV Disinfection Light from UV Can Sanitize Corp. (North Vancouver, BC, Canada) after a minimum 45-minute warm-up time. The lamp’s emission spectrum is tightly peaked around 222 nm (Fig. Figure S2). Before each irradiation, the irradiance (power per unit area, E) at the surface of the sample (1 cm from the lamp) was determined using a Molectron PM5100 Laser Power Meter with a 1 cm^2^ paper mask. From the irradiance, the exposure time (t) necessary to achieve the desired fluence (H) was calculated using the equation H=E×t. For samples in sample chambers, irradiation was performed through a quartz microscope slide with 67% transmittance (as measured using a Molectron PM5100 Laser Power Meter), and exposure times were adjusted accordingly. Exposure times ranged from 4 s to 23 min.

### SDS–PAGE

Samples destined for SDS-PAGE were irradiated in a clear-bottom 96-well plate (see above) with 60 µL per well, corresponding to a sample depth of about 1.9 mm. Each irradiated sample was transferred to a 0.5 mL tube with an equal volume of 2X reducing Laemmli buffer (4% sodium dodecyl sulfate, 10% β-mercaptoethanol, 125 mM Tris–HCl, 20% glycerol, 0.002% bromophenol blue) and heated at 95°C for 10 minutes. Samples were loaded into a discontinuous SDS–PAGE system (5% stacking gel, 6 or 8% resolving gel, 3 or 6 µg protein per well) and run in a Bio-Rad Mini-PROTEAN Tetra Vertical Electrophoresis Cell at 50 mA for approximately 45 minutes alongside a prestained protein ladder (PageRuler Prestained Protein Ladder, 10 to 180 kDa, Thermo Fisher Scientific). Gels were stained for 1 h in a Coomassie solution (0.1% Coomassie brilliant blue R-250, 40% ethanol, 10% acetic acid) and de-stained overnight in 10% ethanol and 7.5% acetic acid. Following de-staining, gels were imaged with an Olympus model C-5060 Wide Zoom camera in a UV transilluminator at 302 nm (High Performance UV Transilluminator, UVP).

### Microscopy

Samples destined for microscopy were irradiated in a 0.175 or 1.2 mm-thick sample chamber (see above). Bright-field microscopy was performed using an Olympus IX83 inverted light microscope with an ORCA-Spark camera (Hamamatsu). Due to the optical resolution limit, only collagen fibrils, not monomeric collagen molecules, are visible in a conventional light microscope. No image post-processing was performed.

## Results and Discussion

Collagen fibrils in 0.175 mm-thick sample chambers were irradiated with 1000, 5000, or 10,000 mJ/cm^2^ of far-UVC radiation. Exposure to 1000 mJ/cm^2^ of far-UVC radiation resulted in significant collagen fibril degradation and disruption of the fibrillar network (Fig. 2A). Exposure to 5000 and 10,000 mJ/cm^2^ of far-UVC radiation caused apparently complete collagen fibril degradation and loss of the fibrillar network.

**Fig 2.**
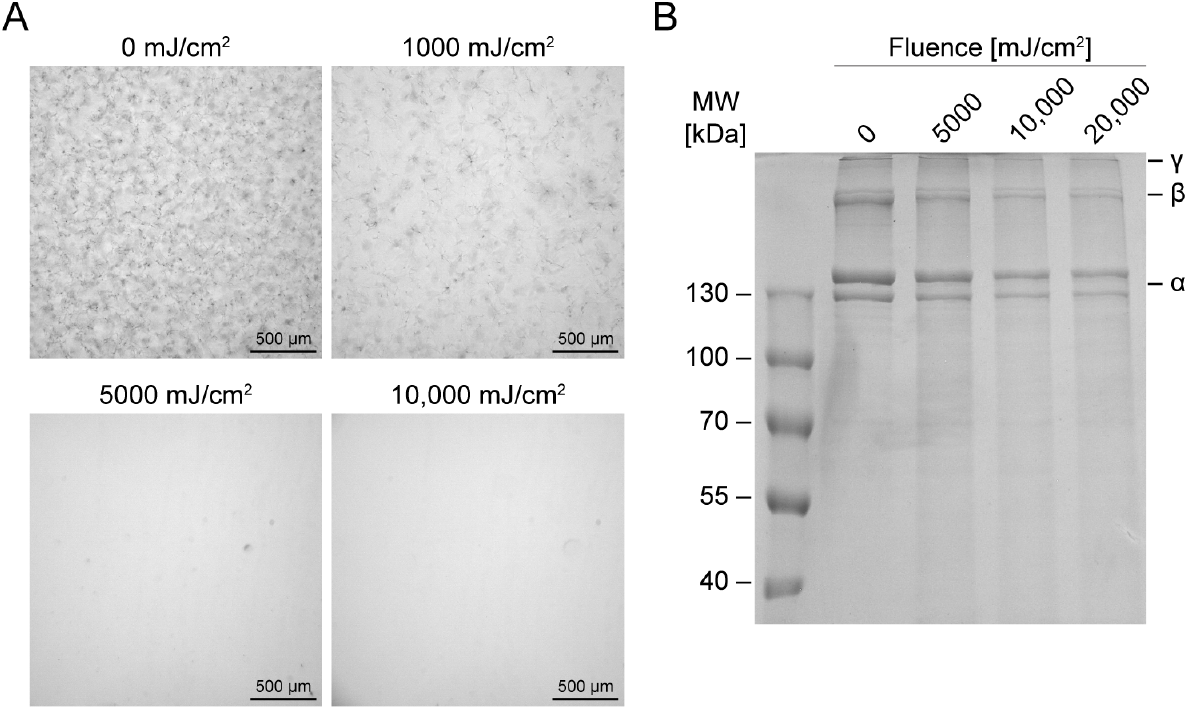
Collagen fibrils before and after irradiation with far-UVC. A: Collagen fibrils exposed to 1000, 5000, and 10,000 mJ/cm^2^ of far-UVC radiation and imaged under a light microscope at 10X magnification. B: Collagen fibrils exposed to 5000, 10,000, and 20,000 mJ/cm^2^ of far-UVC radiation and run on an SDS–PAGE gel (8% resolving gel) with 6 µg protein per well.

To determine whether fibril degradation was a result of depolymerization or chain scission, irradiated fibrils were analyzed by SDS–PAGE. Type I collagen molecules produce five characteristic bands on an SDS–PAGE gel: α_1_ and α_2_ bands with a 2:1 ratio, β_11_ (two cross-linked α_1_ chains) and β_12_ (cross-linked α_1_ and α_2_ chains) bands with a 1:2 ratio, and a γ band (three cross-linked α chains). Following exposures of 5000, 10,000, and 20,000 mJ/cm^2^ of far-UVC radiation, SDS-PAGE analysis revealed a decrease in intensity of collagen α, β, and γ bands with increasing far-UVC fluence (Fig. 2B). This indicates that far-UVC radiation-induced fibril degradation involves polypeptide chain scission. Furthermore, the absence of new low molecular weight bands in the lanes of the irradiated samples suggests that far-UVC radiation-induced chain scission is random and/or results in fragments smaller than 40 kDa. These results are consistent with the existing literature on the effect of 254 nm UV radiation on type I collagen, which reports collagen chain scission at high fluences [33–37]. However, some of the aforementioned studies also describe UV-induced collagen cross-linking, and suggest that the two effects occur simultaneously upon exposure to 254 nm UV radiation [34, 35, 37]. In these cases, cross-linking dominates at low fluences and chain scission at high fluences [34, 35, 37]. Although the SDS-PAGE gel in Figure 2B shows no evidence of cross-linking following far-UVC exposures of 5000 mJ/cm^2^ and above, we are unable to conclude from our data whether low fluences of far-UVC radiation induce collagen cross-linking.

To elucidate whether far-UVC radiation-induced chain scission is dependent on protein structure or sequence, both molecular type I collagen and bovine serum albumin (BSA) were irradiated and subjected to SDS–PAGE. As shown in Figure 3A, there was a greater loss of intensity of collagen α, β, and γ bands for collagen irradiated in molecular form than collagen irradiated with the same fluence in fibrillar form, indicating that the former underwent more chain scission than the latter. However, the optical density at 222 nm of a fibrillar collagen solution is greater than that of a molecular collagen solution due to increased light scattering [31]. Thus, the observed effect may be a function of increased far-UVC attenuation in the fibrillar solution rather than a structure-dependent effect of collagen molecules on the outside of fibrils shielding molecules on the inside. Figure 3B shows that irradiation of BSA with 5000, 10,000, and 20,000 mJ/cm^2^ of far-UVC resulted in a pattern of fluence-dependent band disappearance similar to that observed for fibrillar collagen (Fig. 3B and Fig. 2B). We conclude that exposure to far-UVC radiation can lead to chain scission in both fibrous and globular proteins, and that the fluence-dependent effects are not specific to collagen.

**Fig 3.**
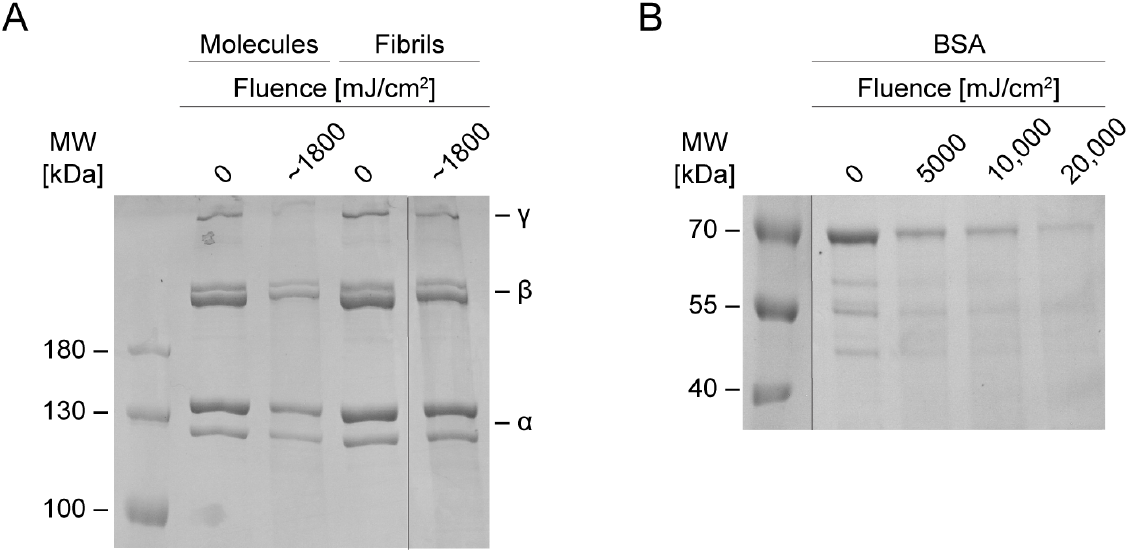
SDS-PAGE gels of molecular collagen, fibrillar collagen, and BSA before and after irradiation with far-UVC. A: Collagen molecules and fibrils exposed to equal fluences of far-UVC radiation and run on a 6% resolving gel with 3 µg protein per well. The black vertical line indicates where two portions of the original gel were spliced together to remove irrelevant lanes. B: BSA molecules irradiated with 5000, 10,000, and 20,000 mJ/cm^2^ of far-UVC radiation and run on an 8% resolving gel with 6 µg protein per well. The black vertical line indicates where two portions of the original gel were spliced together to remove irrelevant lanes.

To further assess how optical density influences fibrillar collagen network disruption by far-UVC radiation, we prepared fibrillar collagen samples with two different thicknesses for irradiation. Upon increasing sample thickness from 0.175 mm to 1.2 mm, we observed a significant reduction in fibril degradation, with intact fibrils present in the 1.2 mm sample even after exposure to approximately 2500 mJ/cm^2^ of far-UVC radiation (Fig. 4). This is consistent with the hypothesis that greater scattering of 222 nm light by the fibrils in the thicker sample leads to stronger radiation attenuation and less overall fibril damage. Translating this to a clinical scenario, if any fibril damage were to occur as a result of far-UVC exposure, it would likely be superficial, restricted to the top layer of tissue.

**Fig 4.**
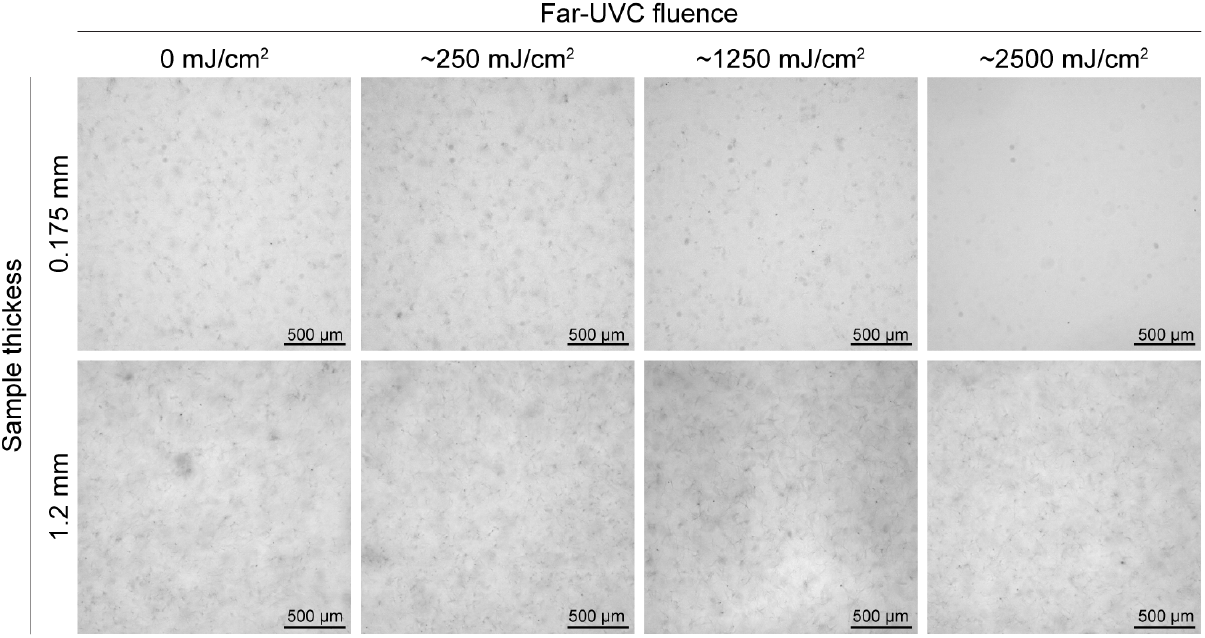
Fibrillar collagen samples of different thicknesses before and after irradiation with far-UVC. Collagen fibrils were irradiated with increasing fluences of far-UVC radiation and imaged under a light microscope at 10X magnification. Top row corresponds to a thin sample, bottom row corresponds to a thick sample.

While these results provide an initial validation of the safety of far-UVC radiation for extracellular matrix and connective tissue exposure, further analysis is needed in more tissue-realistic systems. The collagen fibrils used in this study lack intermolecular cross-links present *in vitro*, as well as non-covalent interactions with cells and other matrix proteins [38]. Furthermore, the fluences employed here are greater than those which would be used clinically. Previous work has established that far-UVC radiation is an effective antibacterial agent at fluences below 100 mJ/cm^2^ [39, 40]. It will be important to investigate whether low fluences of far-UVC radiation induce collagen cross-link formation [34, 35, 37], as increased ECM cross-linking is associated with tissue fibrosis and impaired wound healing [41, 42]. Far-UVC radiation is also ozone-generating, which poses a potential health hazard for both patients and healthcare workers [43]. In our experiments, exposure to far-UVC radiation caused gas bubbles to form within sealed sample chambers, with higher fluences leading to more and larger bubbles (Fig. Figure S3). However, the amount of gas produced was minimal even at a far-UVC fluence of 1000 mJ/cm^2^, 10-fold greater than that needed to inactivate most bacteria.

## Conclusions

In this work, we showed that the response of type I collagen fibrils to far-UVC radiation is dependent on both far-UVC fluence and fibrillar network thickness. At low fluences and high thicknesses—the most representative of a clinical setting—fibrillar collagen is resistant to significant degradation by far-UVC radiation. Only at fluences many times greater than the bactericidal dose did far-UVC radiation induce random polypeptide chain cleavage in both collagen and BSA. We conclude that germicidal far-UVC lamps should continue to be investigated as promising candidates for the prevention of surgical-site infections.

## Supporting information

**Figure S1.**
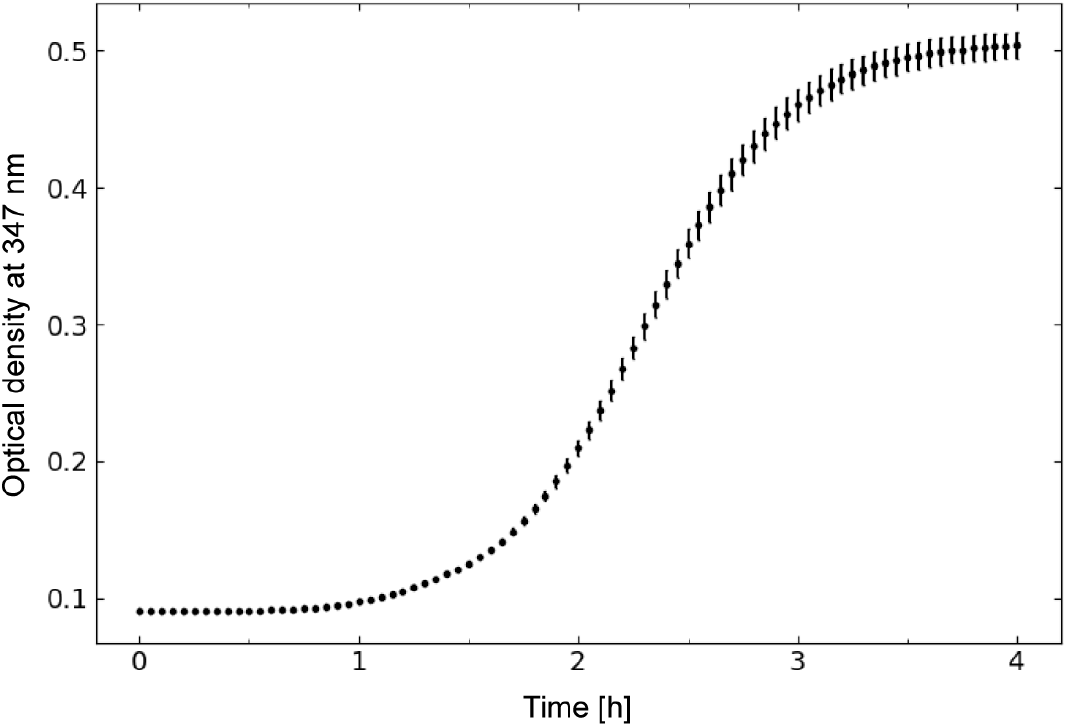
Optical density at 347 nm as a function of time following initiation of collagen fibril formation. Error bars represent the standard error of the mean from three replicate measurements.

**Figure S2.**
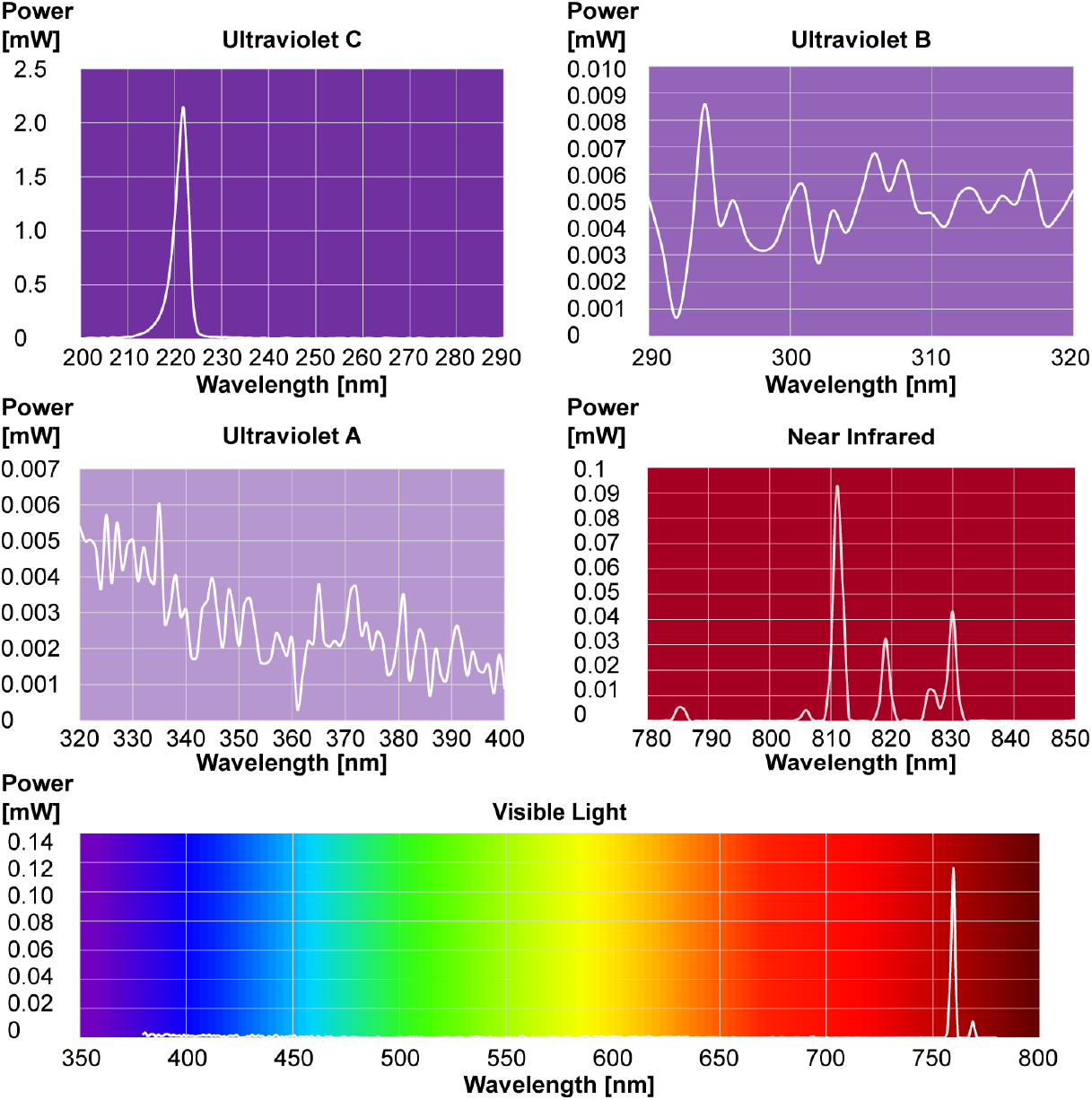
Emission spectra of a LILY Handheld Personal Far UV Disinfection Light. Data courtesy of Viso Systems (Copenhagen, Denmark). Figure adapted from Viso Systems.

**Figure S3.**
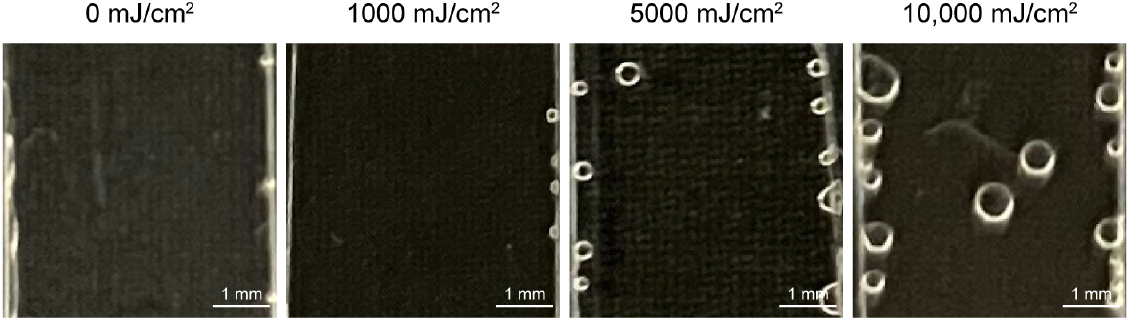
Microscopy sample chambers containing collagen fibrils before and after irradiation with far-UVC. Gas bubbles are increasingly prevalent at higher fluences.

**Figure S4.**
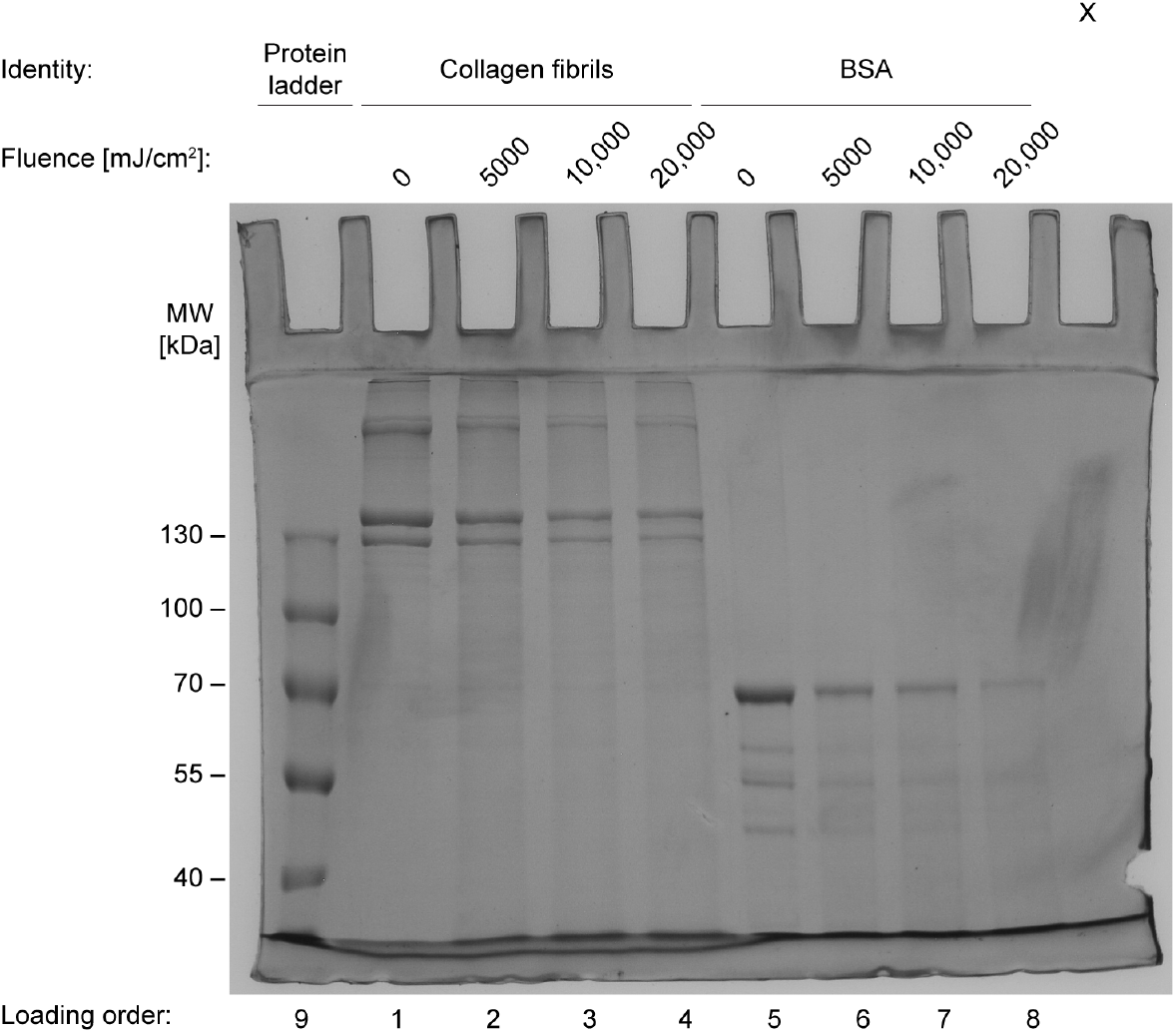
Original SDS-PAGE gel image for Figure 2B and Figure 3B.

**Figure S5.**
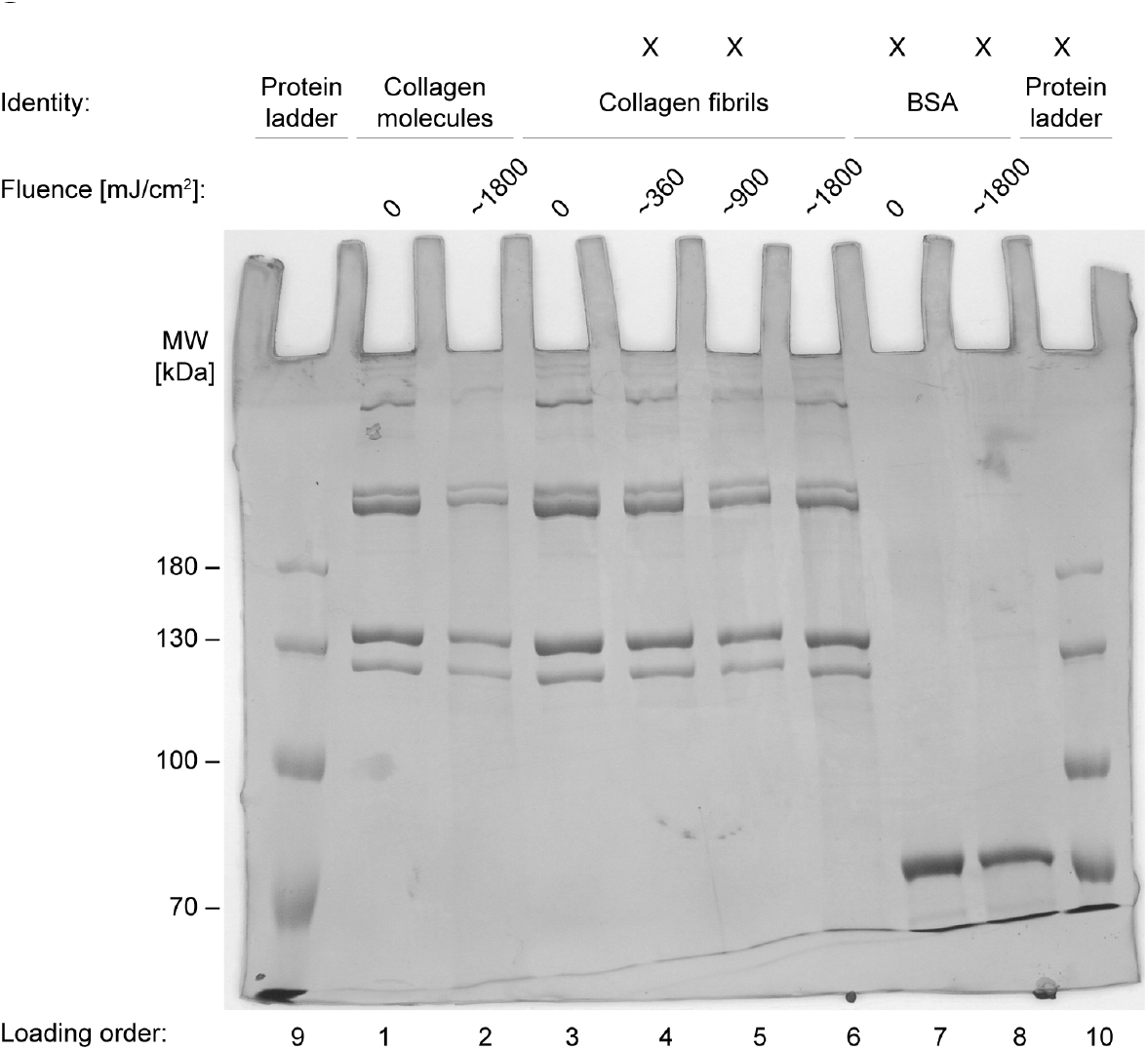
Original SDS-PAGE gel image for Figure 3A.

## Acknowledgments

The authors thank Mehrdad Behnami and Rea-Jayne Smith of UV Can Sanitize Corp. for generously supplying the far-UVC lamp, Jens Lassen and Gary Leach for lending power meters, and Jody Tao and other members of the Forde Lab for their research insights and suggestions. This work was funded by the Natural Sciences and Engineering Research Council of Canada (NSERC) through a Discovery Grant to NRF.

## References

1. World Health Organization. Global guidelines for the prevention of surgical site infection. Geneva: World Health Organization; 2018.

2. Buonanno M, Ponnaiya B, Welch D, Stanislauskas M, Randers-Pehrson G, Smilenov L, et al. Germicidal efficacy and mammalian skin safety of 222-nm UV light. Radiation Research. 2017;187(4). doi:10.1667/RR0010CC.1.

3. Conner-Kerr TA, Sullivan PK, Gaillard J, Franklin ME, Jones RM. The effects of ultraviolet radiation on antibiotic-resistant bacteria in vitro. Ostomy Wound Management. 1998;44(10):50–56.

4. Eadie E, Hiwar W, Fletcher L, Tidswell E, O’Mahoney P, Buonanno M, et al. Far-UVC (222 nm) efficiently inactivates an airborne pathogen in a room-sized chamber. Scientific Reports. 2022;12(1). doi:10.1038/s41598-022-08462-z.

5. Fukui T, Niikura T, Oda T, Kumabe Y, Ohashi H, Sasaki M, et al. Exploratory clinical trial on the safety and bactericidal effect of 222-nm ultraviolet C irradiation in healthy humans. PLOS ONE. 2020;15(8). doi:10.1371/journal.pone.0235948.

6. Goh JC, Fisher D, Hing ECH, Hanjing L, Lin YY, Lim J, et al. Disinfection capabilities of a 222 nm wavelength ultraviolet lighting device: A pilot study. Journal of Wound Care. 2021;30(2). doi:10.12968/jowc.2021.30.2.96.

7. McKinney CW, Pruden A. Ultraviolet disinfection of antibiotic resistant bacteria and their antibiotic resistance genes in water and wastewater. Environmental Science and Technology. 2012;46(24). doi:10.1021/es303652q.

8. Omotani S, Tani K, Aoe M, Esaki S, Nagai K, Hatsuda Y, et al. Bactericidal effects of deep ultraviolet light-emitting diode for solutions during intravenous infusion. International Journal of Medical Sciences. 2018;15(2). doi:10.7150/ijms.22206.

9. Rao BK, Kumar P, Rao S, Gurung B. Bactericidal effect of ultraviolet C (UVC), direct and eiltered through transparent plastic, on gram-positive cocci: An in vitro study. Ostomy Wound Management. 2011;57(7).

10. Taylor W, Camilleri E, Craft DL, Korza G, Granados MR, Peterson J, et al. DNA damage kills bacterial spores and cells exposed to 222-nanometer UV radiation. Applied and Environmental Microbiology. 2020;86(8). doi:10.1128/AEM.03039-19.

11. Welch D, Buonanno M, Shuryak I, Randers-Pehrson G, Spotnitz HM, Brenner DJ. Effect of far ultraviolet light emitted from an optical diffuser on methicillin-resistant Staphylococcus aureus in vitro. PLOS ONE. 2018;13(8). doi:10.1371/journal.pone.0202275.

12. Masschelein WJ, Rice RG. Ultraviolet light in water and wastewater sanitation. Boca Raton: CRC Press; 2002.

13. Pfeifer GP, You YH, Besaratinia A. Mutations induced by ultraviolet light. Mutation Research. 2005;571(1-2):19–31. doi:10.1016/j.mrfmmm.2004.06.057.

14. Narita K, Asano K, Morimoto Y, Igarashi T, Hamblin MR, Dai T, et al. Disinfection and healing effects of 222-nm UVC light on methicillin-resistant Staphylococcus aureus infection in mouse wounds. Journal of Photochemistry and Photobiology B, Biology. 2018;178:10–18. doi:10.1016/j.jphotobiol.2017.10.030.

15. Woods JA, Evans A, Forbes PD, Coates PJ, Gardner J, Valentine RM, et al. The effect of 222-nm UVC phototesting on healthy volunteer skin: A pilot study. Photodermatology Photoimmunology and Photomedicine. 2015;31(3). doi:10.1111/phpp.12156.

16. Buonanno M, Welch D, Shuryak I, Brenner DJ. Far-UVC light (222 nm) efficiently and safely inactivates airborne human coronaviruses. Scientific Reports. 2020;10(1). doi:10.1038/s41598-020-67211-2.

17. Coohill TP. Virus-cell interactions as probes for vacuum-ultraviolet radiation damage and repair. Photochemistry and Photobiology. 1986;44(3). doi:10.1111/j.1751-1097.1986.tb04676.x.

18. Barnard IRM, Eadie E, Wood K. Further evidence that far-UVC for disinfection is unlikely to cause erythema or pre-mutagenic DNA lesions in skin. Photodermatology Photoimmunology and Photomedicine. 2020;36(6). doi:10.1111/phpp.12580.

19. Buonanno M, Randers-Pehrson G, Bigelow AW, Trivedi S, Lowy FD, Spotnitz HM, et al. 207-nm UV light - A promising tool for safe low-cost reduction of surgical site infections. I: In vitro studies. PLOS ONE. 2013;8(10). doi:10.1371/journal.pone.0076968.

20. Buonanno M, Welch D, Brenner DJ. Exposure of human skin models to KrCl excimer lamps: The impact of optical filtering. Photochemistry and Photobiology. 2021;97(3). doi:10.1111/php.13383.

21. Eadie E, Barnard IMR, Ibbotson SH, Wood K. Extreme exposure to filtered far-UVC: A case study. Photochemistry and Photobiology. 2021;97(3). doi:10.1111/php.13385.

22. Hickerson RP, Conneely MJ, Hirata Tsutsumi SK, Wood K, Jackson DN, Ibbotson SH, et al. Minimal, superficial DNA damage in human skin from filtered far-ultraviolet C. British Journal of Dermatology. 2021;184(6). doi:10.1111/bjd.19816.

23. Buonanno M, Stanislauskas M, Ponnaiya B, Bigelow AW, Randers-Pehrson G, Xu Y, et al. 207-nm UV light - A promising tool for safe low-cost reduction of surgical site infections. II: In-vivo safety studies. PLOS ONE. 2016;11(6). doi:10.1371/journal.pone.0138418.

24. Fukui T, Niikura T, Oda T, Kumabe Y, Nishiaki A, Kaigome R, et al. Safety of 222 nm UVC irradiation to the surgical site in a rabbit model. Photochemistry and Photobiology. 2022;98(6). doi:10.1111/php.13620.

25. Lu Y, Zhang S, Wang Y, Ren X, Han J. Molecular mechanisms and clinical manifestations of rare genetic disorders associated with type I collagen. Intractable and Rare Diseases Research. 2019;8(2). doi:10.5582/irdr.2019.01064.

26. Kamrani P, Marston G, Arbor TC, Jan A. Anatomy, Connective Tissue. Treasure Island: StatPearls Publishing; 2022.

27. Shoulders MD, Raines RT. Collagen structure and stability. Annual Review of Biochemistry. 2009;78:929–958. doi:10.1146/annurev.biochem.77.032207.120833.

28. Kirkness MW, Lehmann K, Forde NR. Mechanics and structural stability of the collagen triple helix. Current Opinion in Chemical Biology. 2019;53:98–105. doi:10.1016/j.cbpa.2019.08.001.

29. Li Y, Asadi A, Monroe MR, Douglas EP. pH effects on collagen fibrillogenesis in vitro: Electrostatic interactions and phosphate binding. Materials Science and Engineering C. 2009;29(5). doi:10.1016/j.msec.2009.01.001.

30. Kuznetsova N, Chi SL, Leikin S. Sugars and polyols inhibit fibrillogenesis of type I collagen by disrupting hydrogen-bonded water bridges between the helices. Biochemistry. 1998;37(34). doi:10.1021/bi980089.

31. Williams BR, Gelman RA, Poppke DC, Piez K. Collagen fibril formation. Optimal in vitro conditions and preliminary kinetic results. Journal of Biological Chemistry. 1978;253(18). doi:10.1016/s0021-9258(19)46970-6.

32. Shayegan M, Forde NR. Microrheological Characterization of Collagen Systems: From Molecular Solutions to Fibrillar Gels. PLOS ONE. 2013;8(8). doi:10.1371/journal.pone.0070590.

33. Jariashvili K, Madhan B, Brodsky B, Kuchava A, Namicheishvili L, Metreveli N. UV damage of collagen: Insights from model collagen peptides. Biopolymers. 2012;97(3). doi:10.1002/bip.21725.

34. Miles CA, Sionkowska A, Hulin SL, Sims TJ, Avery NC, Bailey AJ. Identification of an intermediate state in the helix-coil degradation of collagen by ultraviolet light. Journal of Biological Chemistry. 2000;275(42). doi:10.1074/jbc.M002346200.

35. Mori H, Hara M. UV irradiation of Type I collagen gels changed the morphology of the interconnected brain capillary endothelial cells on them. Materials Science and Engineering C. 2020;112. doi:10.1016/j.msec.2020.110907.

36. Rabotyagova OS, Cebe P, Kaplan DL. Collagen structural hierarchy and susceptibility to degradation by ultraviolet radiation. Materials Science and Engineering C. 2008;28(8). doi:10.1016/j.msec.2008.03.012.

37. Sionkowska A, Lewandowska K, Adamiak K. The influence of UV light on rheological properties of collagen extracted from silver carp skin. Materials. 2020;13(19). doi:10.3390/ma13194453.

38. Orgel JPRO, Miller A, Irving TC, Fischetti RF, Hammersley AP, Wess TJ. The in situ supermolecular structure of type I collagen. Structure. 2001;9(11):1061–1069. doi:10.1016/S0969-2126(01)00669-4.

39. Malayeri AH, Mohseni M, Cairns B, Bolton JR. Fluence (UV Dose) Required to Achieve Incremental Log Inactivation of Bacteria, Protozoa, Viruses and Algae. IUVA News. 2016;18(3).

40. Hessling M, Haag R, Sieber N, Vatter P. The impact of far-UVC radiation (200-230 nm) on pathogens, cells, skin, and eyes - a collection and analysis of a hundred years of data. GMS Hygiene and Infection Control. 2021;16. doi:10.3205/dgkh000378.

41. Ewald CY. The matrisome during aging and longevity: A systems-level approach toward defining matreotypes promoting healthy aging. Gerontology. 2020;66(3). doi:10.1159/000504295.

42. Kong W, Lyu C, Liao H, Du Y. Collagen crosslinking: Effect on structure, mechanics and fibrosis progression. Biomedical Materials. 2021;16(6). doi:10.1088/1748-605X/ac2b79.

43. Peng Z, Day DA, Symonds G, Jenks O, Stark H, Anne V, et al. Significant production of ozone from germicidal UV lights at 222 nm. Environmental Science & Technology Letters. 2023;10(8):668–674. doi:10.1021/acs.estlett.3c00314.

